# Adaptive conformational restraints for interactive model rebuilding in Cartesian and torsion space

**DOI:** 10.1101/2020.09.29.318543

**Authors:** Tristan I Croll, Randy J Read

**Affiliations:** Cambridge Institute for Medical Research, Keith Peters Building, Cambridge, Cambridgeshire, CB2 0XY, United Kingdom

**Keywords:** Model building, restraints, refinement

## Abstract

When building atomic models into weak and/or low-resolution density, a common strategy is to restrain its conformation to that of a higher-resolution model of the same or similar sequence. When doing so, it is important to avoid over-restraining to the reference model in the face of disagreement with the experimental data. The most common strategy for this is the use of “top-out” potentials. These act like simple harmonic restraints within a defined range, but gradually weaken when the deviation between the model and reference grows larger than a defined transition point. In each current implementation, the rate at which the potential flattens beyond the transition region follows a fixed form – although the form chosen varies between implementations. A restraint potential with a tuneable rate of flattening would provide greater flexibility to encode the confidence in any given restraint. Here we describe two new such potentials: a Cartesian distance restraint derived from a recent generalisation of common loss functions, and a periodic torsion restraint based on a renormalisation of the von Mises distribution. Further, we describe their implementation as user-adjustable/switchable restraints in ISOLDE, and demonstrate their use in some real-world examples.

**Synopsis:** New forms of adaptive or top-out distance and torsion restraints are described, suitable for restraining a model to match a reference structure during interactive rebuilding. In addition, their implementation in ISOLDE is described along with some illustrative example applications.

## 1. Introduction

Refinement of low-resolution macromolecular models is often an underdetermined problem - that is, even accounting for the restraints imposed by atomic bond, angle and torsion geometry there remain more tuneable parameters than experimental observations. As such, without the imposition of further restraints refinement results become increasingly poor as resolutions degrade beyond 2.5-3Å. While limiting the degrees of freedom by constraining all bond lengths and angles to ideal values (Schröder *et al*., 2010) or the inclusion of explicit van der Waals and electrostatic terms (Croll, 2018; Moriarty *et al*., 2020) can extend the resolutions at which (given a reasonable starting model) good results can be obtained to the high-3Å or low-4Å range, at lower resolutions the most sensible approach is often to take advantage of the information available from higher-resolution structures of similar proteins. This may take the form of restraints on matching torsions as used in *Phenix* (Headd *et al*., 2012) or interatomic distances as used in *REFMAC5* and *Coot* (Nicholls *et al*., 2012) via *ProSMART*(Nicholls *et al*., 2014). Such restraints are implemented as so-called “top-out” potentials – that is, their penalty functions begin to flatten out (and hence impose a progressively weaker bias towards the template) once the deviation between model and template becomes too great, with the intent of allowing real deviations supported by the data while restraining regions where the model, template and data agree.

To date, these restraint schemes have typically been limited in terms of the form of the fall-off at large deviations: while the potential close to the target is proportional to deviation squared in all cases, *ProSMART* uses the Geman-McClure function whereby the long-range potential is proportional to the square root of the deviation; *Phenix* uses the Welsch robust estimator function which flattens to a constant. For the sake of clarity, these forms correspond to a long-range biasing force which is inversely proportional to the deviation (*ProSMART*) or zero (*Phenix*).

Recently, a more general penalty function was described (Barron, 2019) which allows the rate of fall-off (conceptually related to the level of confidence in a given restraint) to itself become a tuneable parameter. This appears to hold significant promise for use in the macromolecular refinement space, where the best reference model(s) may be of only modest homology, in different conformations, or themselves contain modelling errors. Here we describe the extension of this function to include a “flat bottom” tolerance region around the target for the imposition of distance restraints similar to *ProSMART*, and further derive a periodic torsion restraint potential with similar properties. In addition, we demonstrate their implementation in *ISOLDE* (Croll, 2018) and their application in some illustrative examples.

## 2. Restraint derivations

### 2.1. Adaptive distance restraints

Distance restraint potentials were derived based upon the generalised loss function described in (Barron, 2019), modified to include a flat bottom (that is, a range of distances close to the target for which no force is applied). The restraint potential is defined as:

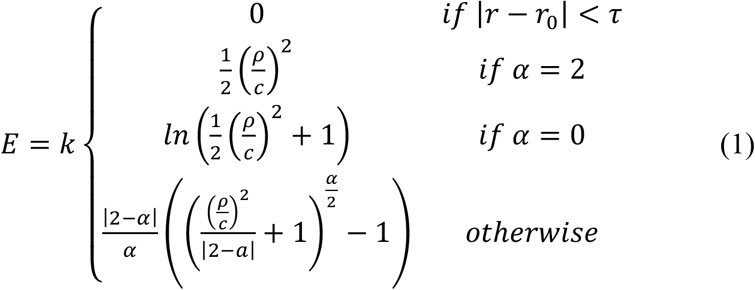

where

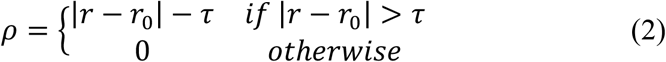

Here, *k* is a spring constant, *r* and *r_0_* are the current and target distances between two restrained atoms respectively; *c* defines the width of the region where the restraint remains approximately quadratic; *α* defines the rate at which the potential flattens outside of the quadratic region, and *τ* is the allowed deviation from *r_0_* for which no penalty is applied.

### 2.2. Adaptive torsion restraints

Since the difference between two angular values θ-θ_0_ is an inherently periodic function, it is sensible for the restraining potential to itself take a periodic form. While non-periodic restraining potentials are typically well-behaved if their gradient is close to zero when θ-θ_0_ = ±180° (that is, when the width of the “well” around the target is small), any non-zero gradient here yields a sharp discontinuity in the first and second derivatives with the subsequent potential for numerical instability. To our knowledge, a periodic penalty function has not previously been described.

In order to develop a suitable potential, we began with the von Mises distribution (Figure 2), a periodic analogue to the normal distribution:

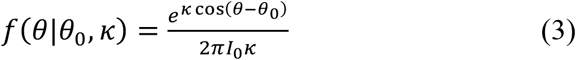

where κ is a shape parameter analogous to the reciprocal of the variance of a normal distribution and *I_0_* is the modified Bessel function of order 0. While this follows the general form required for a periodic top-out potential, it has the undesirable feature that its strength (i.e. the maximum gradient) is a non-obvious function of *κ*, becoming infinitely weak as *κ*, approaches zero (equivalent to expanding the width of the well to its maximum ±180°). Arguably, it is more ideal for a top-out potential to take a form such that the *strength* of the restraint is independent of the *width* of its effective well. To achieve this, we undertook a renormalisation of the von Mises distribution such that the absolute value of its maximum gradient is always 1.

**Figure 1.**
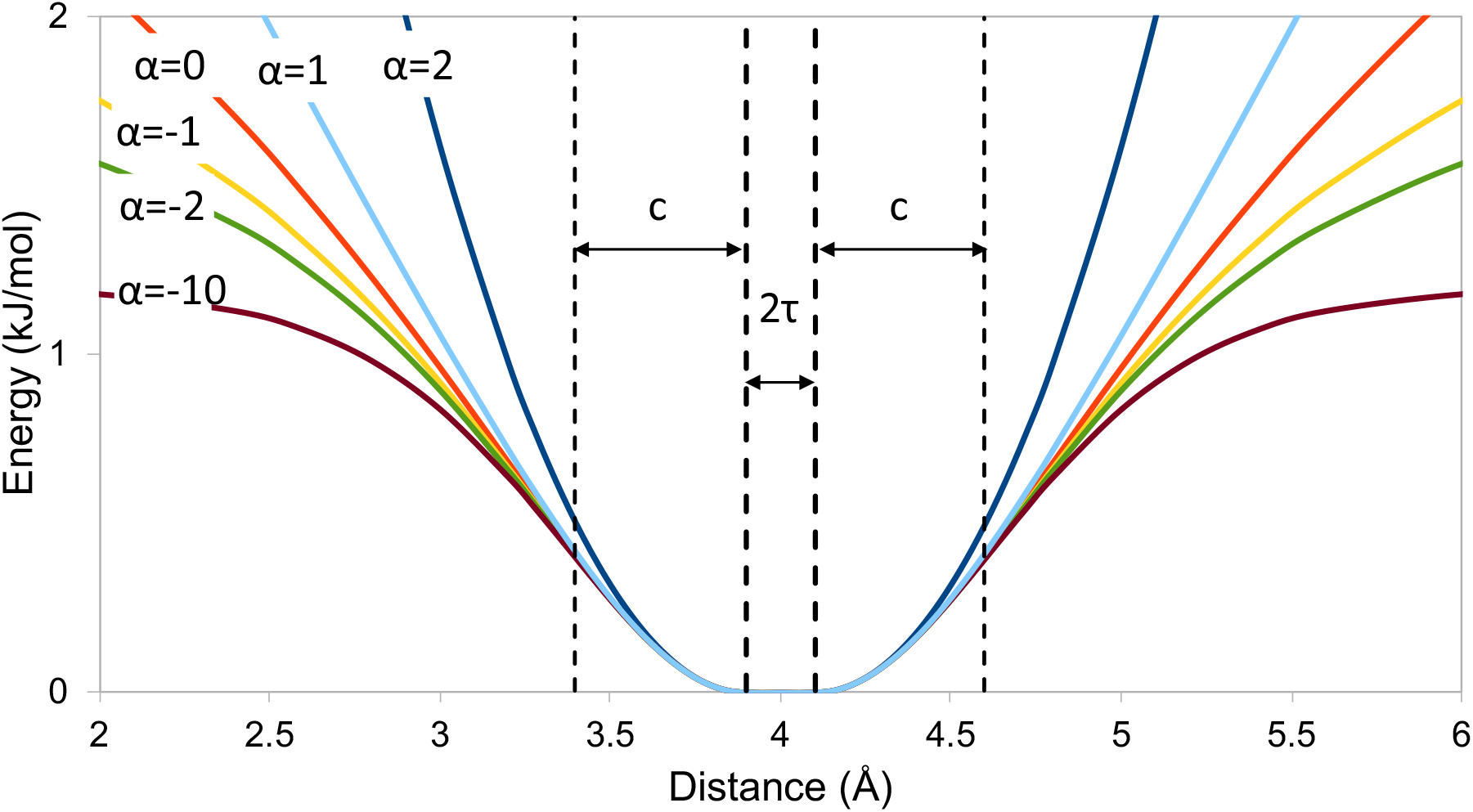
Adaptive distance restraint potential, with the parameters *r_0_* = 4, *τ* = 0.1, *c*=0.5, *k*=1.

**Figure 2.**
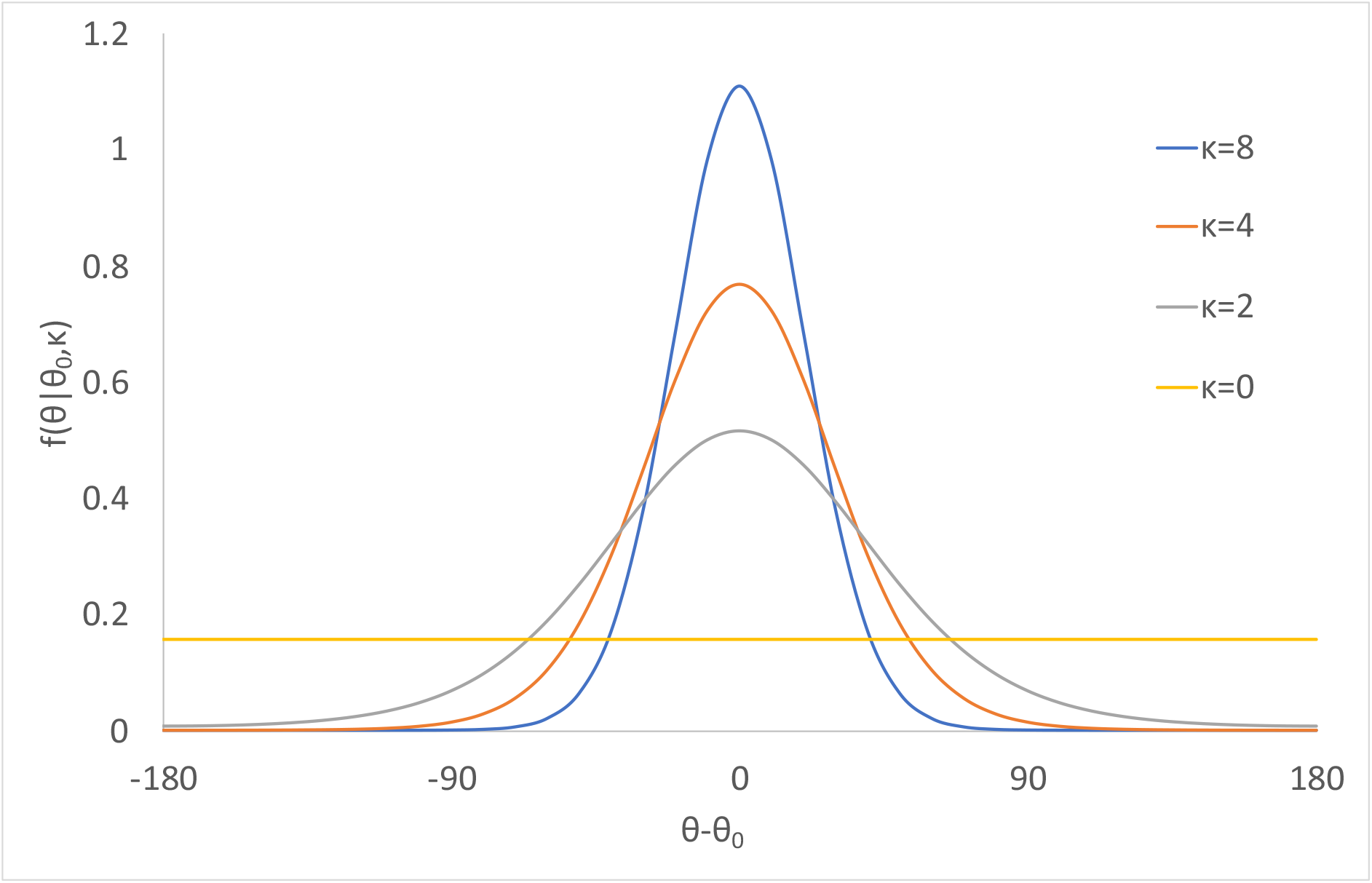
The von Mises distribution. While this has the general form necessary for a periodic top-out potential, it is normalised such that the area under the curve is always equal to 1. The undesirable outcome of this is that the steepness of the well is dependent on its width, and tends to a flat line as κ approaches zero.

Given that a penalty function should reach its minimum when the deviation from the target is zero, we take our starting point as the negative of the numerator of the von Mises distribution:

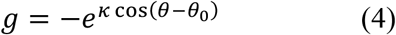

Then:

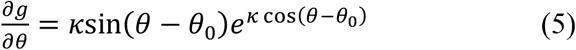

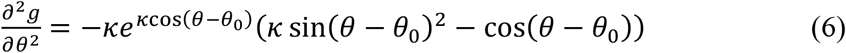

Solving for 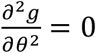 shows that 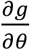 reaches a maximum when:

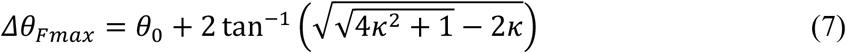

Substituting this into (5) and simplifying yields:

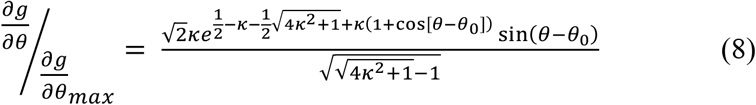

Integrating with respect to θ yields:

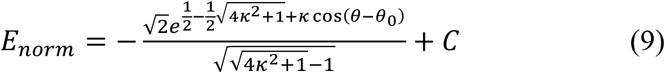

where *C* is a constant of integration. While this is somewhat arbitrary given that the applied bias depends only on the derivative of the potential, it is convenient to set its value to 1 – *E*_*norm*|*C*=0, *θ*–*θ*_0_=*π*_, yielding the form shown in Figure 3:

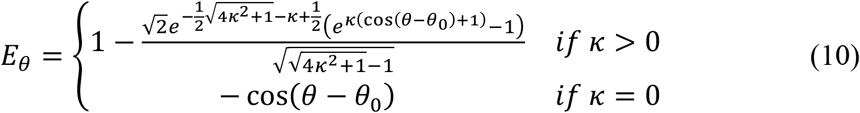

**Figure 3.**
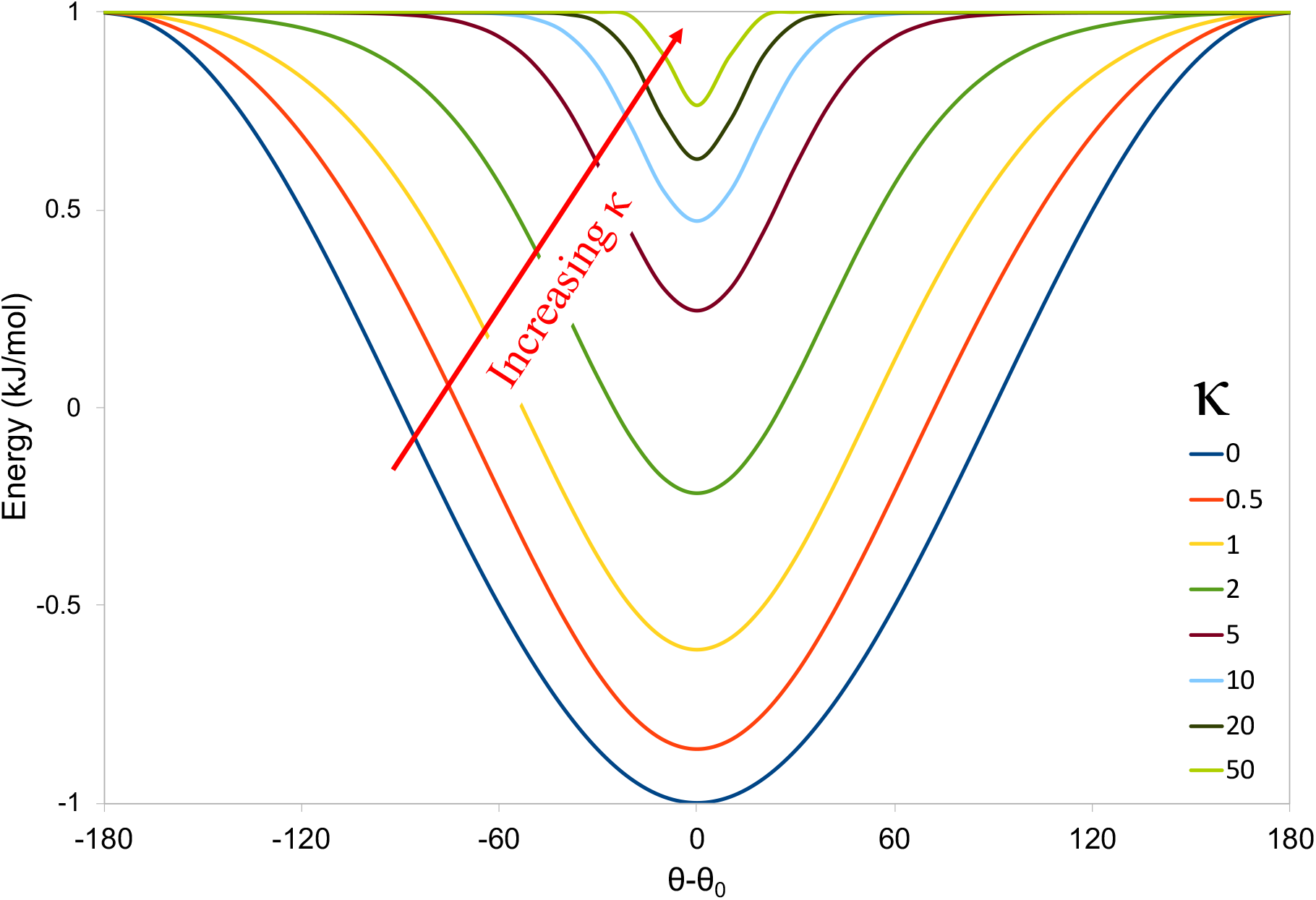
Top-out torsion restraint potential defined in (10).

**Figure 4.**
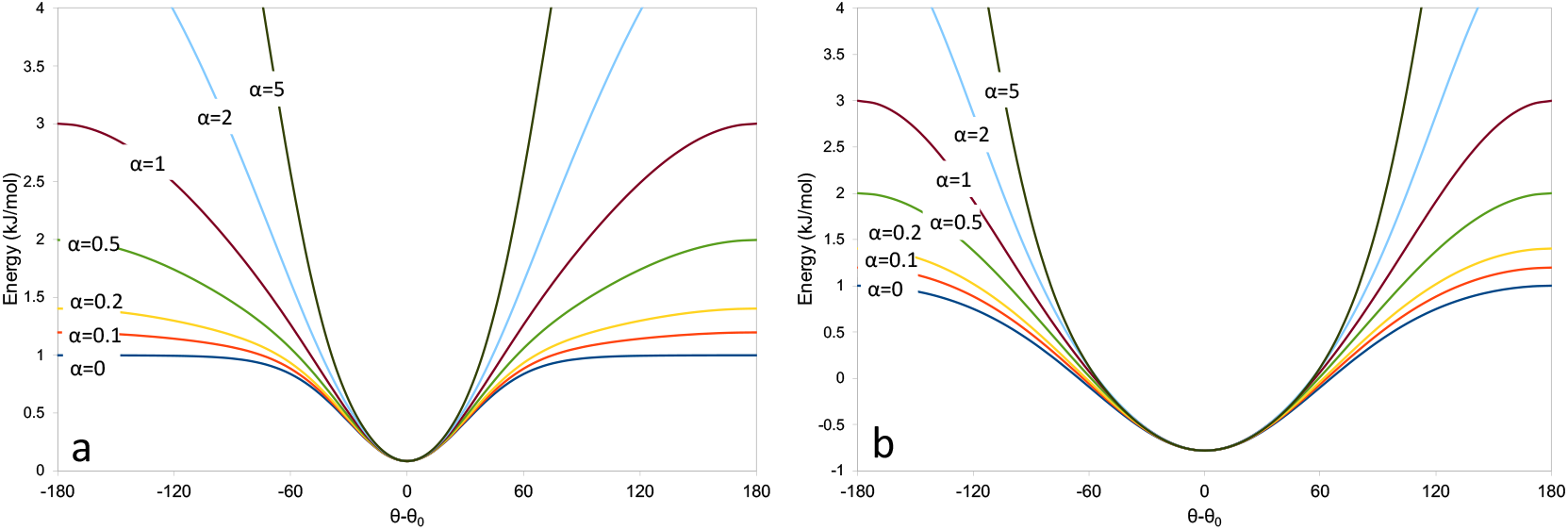
Adaptive torsion restraint potential (12) for (a) Δθ_0_=60° (κ=3.46) or (b) Δθ_0_=120° (κ=0.67).

A more natural definition than κ for the width of the energy well is the value of θ-θ_0_ at which the applied force drops to near zero, defined here as 2Δθ*_F_max__* (equivalent to two standard deviations for small values of θ-θ_0_). If we define this as Δθ_0_, then:

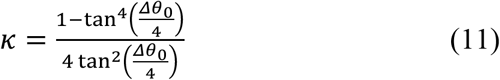

While this potential function displays substantial utility as-is (as will be shown below), it has the remaining drawback that outside of the well region the potential is essentially flat. This is less flexible than the distance-based potential (1), for which the rate of fall-off outside the well is itself a tuneable parameter. If we take *E_θ_* as defined in (10), a potential with tuneable fall-off parameter *α* may be defined as:

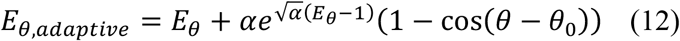

While in principle α is unbounded, in practice values between 0 and 0.5 appear most useful. Negative values cause the potential outside the well to become repulsive; values larger than 1 lead to restraints that are steeper than a straightforward cosine. When α=0 the potential is identical to (10). It is important to note that in contrast to (10) the maximum gradient is no longer strictly independent of *κ* for non-zero values of α, but the variation is small (typically 20-40%) for 0 ≤ α ≤ 0.5.

## 3. Implementations

The adaptive distance and torsion restraints are implemented in ISOLDE (Croll, 2018) using the `CustomBondForce` and `CustomTorsionForce` classes in OpenMM (Eastman *et al*., 2017) and exposed to the user via the ChimeraX command line (Pettersen *et al*., 2020) as the commands `isolde restrain distances` and `isolde restrain torsions` respectively. In each case, the choice is provided to restrain the model to its current geometry or that of a homologous template.

### 3.1. Isolde restrain distances command

Various options are provided for restraining the model either to its own coordinates or to a homologous template. In the most general case, a selection of chains (or fragments thereof) is restrained to a matched selection from the template. The general form of the necessary command is:

~~~
isolde restrain distances {model selection} template
{reference selection}
~~~

The model and reference selections should be matched, comma-separated lists of subselections, where each sub-selection encompasses residues from a single chain. For example, to restrain a fragment consisting of all of chain A and residues 50-100 of chain B from the working model to match the corresponding portion of chains C and D from a template model, the selections would be `#1/A,#1/B:50-100` and `#2/C,#2/D` respectively. It is usually not necessary to specify an explicit range of residues in the template selection – these will be automatically assigned by sequence alignment as described below. An optional argument, `perChain`, is used to decide whether interfaces between chains are restrained (true by default). Note that the template selection need not come from a different model – restraining to the geometry of other chains within the same model is also supported. Restraints are applied using the following protocol:

1. All protein and nucleic acid residues defined by the first selection are concatenated into a single super-sequence, and the same is done for the template selection.
2. These two sequences are then aligned using a secondary structure matching algorithm (implemented as part of the ChimeraX *MatchMaker* tool) to give a list of paired atoms, where each atom is the “principal” atom from its residue (CA for protein, C4’ for nucleic acids). Residues which cannot be matched at this step will not be restrained.
3. The paired sets of atoms are then aligned to find the largest pseudo-rigid body within which all atoms differ in position by less than a tolerance set by the `alignmentCutoff` argument (5Å by default).
4. Residues whose principal atoms fall within the alignment at step 3 are restrained as follows:
  a. A list of paired atoms is generated (atoms with names from Table 1 that appear in both paired residues).
  b. For each atom pair, all other template atoms in the list (excluding those from the same residue) coming within a cutoff distance (default 8Å, adjustable via the optional `distanceCutoff` argument) of the current template atom are found.
  c. For each found template atom, a corresponding restraint is set in the model according to equation (1) with target distance *r_0_* equal to the distance seen in the template. Each restraint is given a flat bottom by default, with τ=0.25*r_0_* (adjustable via the optional `tolerance` argument). The flattening parameter α is set to −2-`fallOff` *\*ln*(*r_0_*) based on the reasoning that larger distances are inherently less certain. For the same reason, the half-width of the harmonic well, *c*, is proportional to the template distance (*c*=0.05*r_0_* by default, adjustable via the `wellHalfWidth` parameter).
5. Steps 3-4 are repeated for any residues not captured by the previous rigid-body alignment, and iterated until it becomes impossible to align at least 3 residues. This allows reasonable restraints to still be applied when the relative orientation of domains is different between model and template.

**Table 1.** Default atoms restrained with adaptive distance restraints in ISOLDE. Since the number of restraints increases geometrically with the number of different atom types included this list is kept small, relying on the molecular dynamics force field to maintain the geometry of the remaining atoms. Other atoms may also be restrained using the `customAtomNames` argument to `isolde restrain distances`. If necessary, these may be combined with torsional restraints as described below.

| Residue type | Restrained atoms |
| --- | --- |
| Protein | CA, CB, CG, CG1, OG, OG1 |
| Nucleic acid | OP1, OP2, C4', C2', O2, O4, N4, N2, O6, N1, N6 |

While restraining to a separate template model is supported as described above, it is likely that the most common use of these restraints will be in restraining the working model to its own starting coordinates prior to undertaking large-scale bulk rearrangements – e.g. when refitting an existing model into a new cryo-EM map of the same complex in a different conformation. An example of this is provided in ISOLDE as a tutorial (accessible via the `isolde tut` command), and involves refitting a model of the ATP-bound state of the *E. coli* LptB2FG transporter (PDB ID 6mhz) into the map associated with its ATP-free state (PDB ID 6mhu; EMDB ID 9118) (Li *et al*., 2019). Figure 5 shows the interface between a pair of helices adjacent to the ATP binding site following refitting. This interface opens substantially in the ATP-free state; the restraints shown in purple have stretched beyond the harmonic well due to the concerted influence of the map and local atomic interactions. In such situations where a subset of restraints clearly disagree with the map it is sensible to selectively release them; this can be achieved using the `isolde release distances` command.

**Figure 5.**
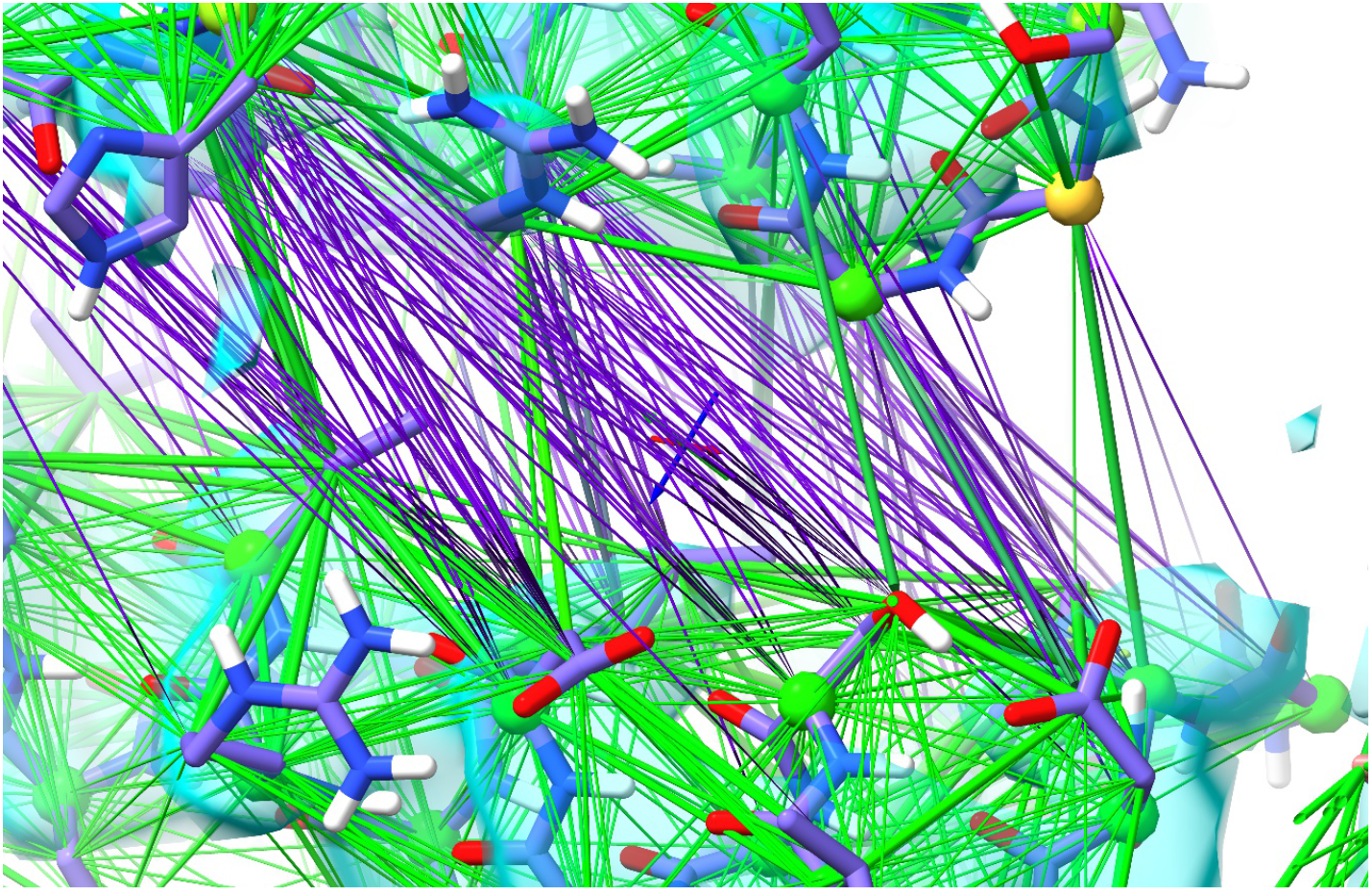
Adaptive distance restraints in ISOLDE around the ATP-binding site of 6mhz after refitting into the map corresponding to the ATP-free state 6mhu. Each restraint is represented as a cylinder whose thickness corresponds to the applied force. Restraints in purple have been stretched beyond the harmonic region causing their applied force to reduce. This is further demonstrated in Supplementary Movie 1.

### 3.2. Isolde restrain torsions command

Restraining the torsions in one protein chain (or selection thereof) to another is achieved in ISOLDE using a command of the form:

~~~
isolde restrain torsions {model selection} template {reference selection}
~~~

where the model and reference selections are each from a single chain. The model may also be restrained to its current conformation by omitting the `template` argument – in this case the model selection may encompass multiple chains. These restraints are currently only supported for protein residues. The parameters of each applied restraint may be modified using the optional arguments `angleRange` (default 30°) to adjust the width of the well, `springConstant` (default 250 kJ/mol) to set the strength of the restraints, and alpha (default 0.2) to set the falloff rate. Additionally, the `backbone` and `sidechains` arguments (both true by default) specify whether restraints should be applied to backbone or sidechain torsions.

In order to assign the restraints, the model and reference sequences are first aligned using the same algorithm as for the adaptive distance restraints. Residues that do not align are not restrained. By default, sidechain torsions are only restrained for identical residues. Peptide bond ω dihedrals are not restrained with adaptive restraints; instead a cosine potential with a ±30° flat bottom is used to restrain them to *cis* or *trans* according to the reference model, with the exception that sites that are *cis*-proline in the template but non-proline in the model will be left in their original conformation.

An example of the depiction of these restraints is shown in Figure 6.

**Figure 6.**
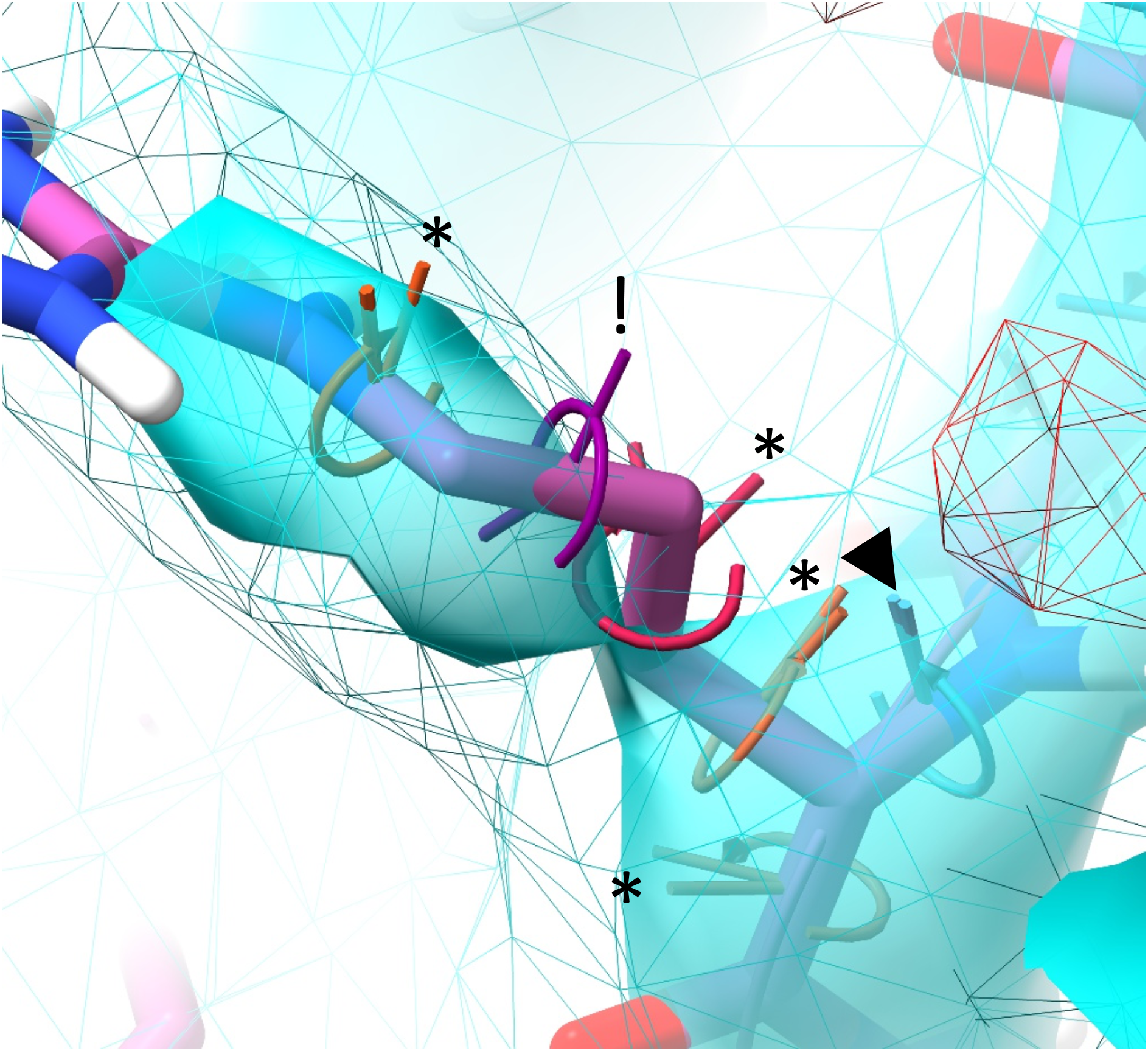
Adaptive torsion restraints with an angle range of 120° applied to an arginine residue. Satisfied restraints (▼) are coloured cyan; the colour shades through orange to red for unsatisfied restraints that are within the restraint well (*); restraints for which the current torsion is outside the well (!) are coloured purple. The angle between the two “posts” indicates the current deviation between the torsion and the restraint target.

## 4. Effect of torsion restraint parameters

As a testbed for the effects of the `angleRange` and `alpha` parameters we chose 3fyj (Anderson *et al*., 2009). This 3.8Å, 282-residue structure of MAPKAP kinase-2 appears to have received only preliminary refinement prior to deposition, and as such appears to be a reasonable facsimile of a modern early-stage model. While higher-resolution crystals of the same protein exist, in order to generate a more realistic scenario we chose as our reference model the 74% identical, 1.8Å model of MAPKAP kinase-3, 3fhr (Cheng *et al*., 2010). In order to obtain the best possible high-resolution reference model, we first did one round of rebuilding and refinement of 3fhr. Manual checking and (where necessary) rebuilding of the reference model is often advisable, particularly for older models; in many cases the output from automatic rebuilding and re-refinement by PDB_REDO (Joosten *et al*., 2014) may be a better starting point than one downloaded directly from the wwPDB. Additionally, as an extra point of comparison we performed a thorough rebuild and re-refinement of 3fyj, with three rounds of end-to-end inspection/correction in ISOLDE interspersed with restrained refinement in *Phenix* beginning from a model settled with `angleRange`=120°, `alpha`=0. Before-and-after validation statistics for both crystals are shown in Table 2.

**Table 2.** Validation statistics for models rebuilt in this work.

|  | 3fyj | Rebuilt | 3fhr | Rebuilt |
| --- | --- | --- | --- | --- |
| Resolution | 3.8Å | 3.8Å | 1.8Å | 1.8Å |
| R<sub>work</sub> | 0.328 (0.265)* | 0.234 | 0.226 | 0.212 |
| R<sub>free</sub> | 0.388 (0.317)* | 0.275 | 0.267 | 0.236 |
| Ramachandran outliers (%) | 14.86 | 0.00 | 0.00 | 0.00 |
| Favoured (%) | 56.52 | 96.39 | 95.85 | 97.36 |
| Ramachandran Z-score | -6.81 | -0.4 | -2.05 | -0.37 |
| Rotamer outliers | 20.31 | 0.00 | 8.13 | 0.00 |
| Clashscore | 69.56 | 2.35 | 7.87 | 3.82 |
| CaBLAM outliers (%) | 13.3 | 1.1 | 0.8 | 0.4 |
| MolProbity score | 4.24 | 1.26 | 2.41 | 1.29 |

\* 3fyj was originally refined with a single overall B-factor, leading to very high R-factors. The results after B-factor-only refinement in *phenix.refine* are shown in parentheses.

For each of the scenarios summarised in Figure 7, the same essential protocol was followed. In brief, the original 3fyj model was restrained to the torsions of the rebuilt 3fhr with the desired parameters, and settled in ISOLDE with temperature gradually reduced from 100-0K over the course of approximately 1 minute (about 50,000 simulation timesteps). An example of this is shown in Supplementary Movie 2. The resulting coordinates were then refined in *phenix.refine* (6 refinement rounds of reciprocal-space XYZ and individual B-factor refinement, using the starting coordinates as a reference model). To define “incorrect” residues, we compared each model to our manually rebuilt and refined exemplar, using a backbone and sidechain torsion scoring system we defined for assessment of CASP13 model predictions (Croll *et al*., 2019) (for backbones the average of unit chord lengths arising from Δφ, Δψ and Δω; for sidechains a weighted average of Δχ_1_ and Δχ_2_ chord lengths adjusted for the degree to which the sidechain is buried). In each case an “incorrect” residue was defined as one with a score higher than 0.15 (approximately equivalent to an average deviation of ±45° from the exemplar). Using only the top-out potential (10) (panels a-c) we saw a broad optimum in R-factors and MolProbity score for 90°`≤angleRange≤`150°. While a slight reduction in the number of incorrect sidechains was observed for `angleRange`=180° (that is, a pure cosine potential), this came at the expense of significant increases in both R-factors and MolProbity score.

**Figure 7.**
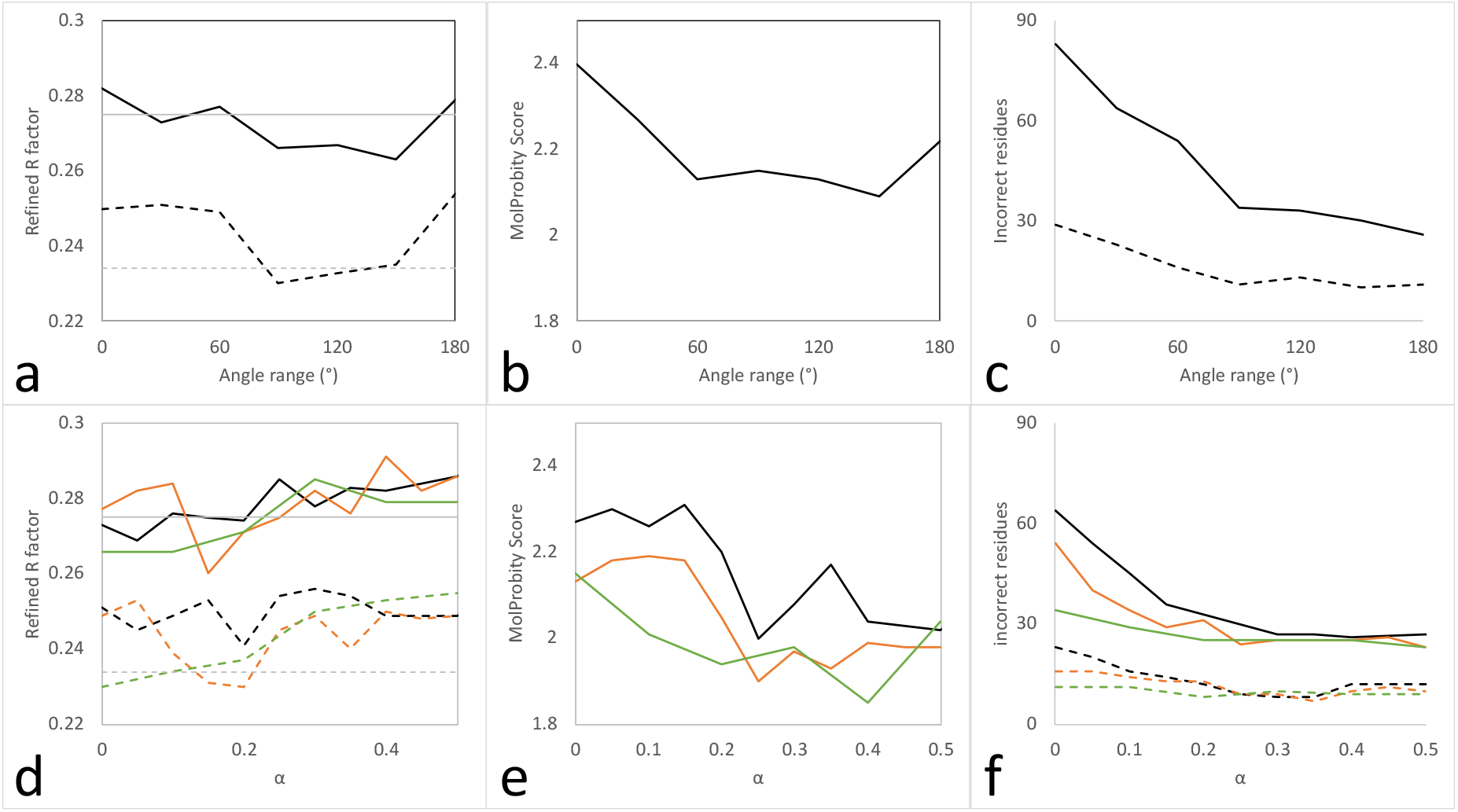
Effect of top-out (10) or adaptive (12) torsion restraints on refinement of 3fyj using an optimised 3fhr as a reference. All models were settled in ISOLDE for ~1 min then refined in *Phenix* as described in the main text. The restraint spring constant was fixed at 250 kJ/mol. (a-c) Top-out restraints only; (d-f) adaptive restraints, angle range=30° (black), 60° (orange) or 90° (green). (a,d) Refined R_free_ (solid lines) and R_work_ (dashed lines). R-factors for the fully rebuilt and refined exemplar are shown as grey horizontal lines. (b,e) MolProbity score (1.29 for the exemplar model). (c, f) Number of remaining large sidechain (solid lines) or backbone (dashed lines) deviations from the exemplar model.

Introduction of the `alpha` parameter (panels d-f) yielded clear improvements over the pure top-out potential for smaller `angleRange` values. For 30°`≤angleRange≤`90° the most reasonable value of `alpha` appeared to be around 0.2: higher values yielded little further improvement in the model geometry and increased the R-factors. The defaults in the current release of ISOLDE have been set to `angleRange`=60° and `alpha`=0.2.

## 5. Discussion

When considering the application of top-out restraints, it is important to note that the requirements of an interactive model-building environment are subtly different from those of non-interactive refinement. In the latter situation, since the results are typically not thoroughly inspected until the (often long-running) refinement process is complete the aim is generally to first do no harm. That is, it is generally preferable to err towards restraints with a small harmonic region to avoid overly-aggressive forcing of the model to the template conformation at the expense of the data. Thus, the default settings in *phenix.refine* only impose strong restraints to model torsions within about ±30° of their counterpart in the template; in the *ProSMART-REFMAC5* pipeline in *CCP4i2* (Potterton *et al*., 2018) restraints fall off beyond ±0.15Å from the target distance and are only applied to distances <4.5Å.

In an interactive environment, on the other hand, the impact of “over-zealous” restraints is arguably less serious since the user is able to immediately observe their local effects in context with the experimental density and may then choose to (selectively) adjust or release them or reset the model to the pre-restrained state and try again. In this context, it becomes much more important to emphasize stability over a wide range of parameter values and initial deviations from the target to provide as much flexibility to the practitioner as possible. Given that the most common use we envision for these restraints in ISOLDE will be to quickly improve a preliminary model (*e.g*. one derived from an auto-building program), we have set the default parameters to be somewhat broader than their analogues in *Phenix* and *REFMAC5:* the torsion restraint well is ±60° with a non-zero gradient beyond that point; distance restraints are applied to interatomic distances <8Å and have a wider well than in *REFMAC5* for all distances >3Å (albeit with a faster fall-off outside the well).

It is important to emphasize that these restraints (and reference-model restraints in general) should be seen as an adjunct to, rather than replacement for, manual inspection and rebuilding. As seen in Figure 7, after settling and refining with optimised torsion restraints around 30 of 282 residues remained significantly different from the model obtained by extensive rebuilding; while many of these arose simply due to the fact that their identity differed between model and template (and hence were unrestrained in their sidechains), others were due to fundamental local conformational differences where the starting conformation was nevertheless close enough to fall into the restraint well, or sites where the model and template *should* match but were too dissimilar in conformation for the restraints to take effect. In such situations direct human intervention remains the safest approach. The visualisations in ISOLDE are designed to make unsatisfied restraints immediately apparent by eye; a future tool will also list these to support systematic inspection.

Finally, we note that all current implementations of top-out or adaptive restraints in the context of macromolecular model building (including those described here) do not take full advantage of their potential. Currently, the parameters controlling the restraint shape and strength are either global to all restraints or (in the case of our distance restraint implementation) simple functions of distance. However, this need not be the case. The precise form of each individual should be set via a Bayesian strategy - based upon confidence regarding our prior information for that particular site. A non-exhaustive list of inputs to such an approach may include conservation in multiple sequence alignment, agreement in multiple structure alignment, correlated conformations for conservatively-substituted residues, local conformational flexibility (estimated via structure alignment and/or local B-factor relative to the bulk), or degree of solvent exposure. This will be an avenue of research for further work.

## Acknowledgements

We gratefully thank Dr Airlie McCoy for her helpful comments and suggestions during the drafting of this manuscript. This work was supported by funding from Wellcome Trust grant 209407/Z/17/Z.

## Supporting information

Supplementary Movie 1: Example use of adaptive distance restraints in refitting a model into a cryo-EM map of substantially different conformation. An interactive tutorial covering this scenario is available in ISOLDE via the command `isolde tut`.

Supplementary Movie 2: Use of adaptive distance restraints to restrain 3fhr to a template model obtained by rebuilding the higher-resolution 3fyf, and initial subsequent steps of manual inspection and rebuilding.

